# Compact longitudinal representations derived from mixed-format lifestyle questionnaires outperform static text-derived features for ALS-versus-control classification

**DOI:** 10.64898/2026.03.23.713709

**Authors:** Jorge Radlowski Nova, Juan I. Lopez-Carbonero, Fernando J. Gómez-Márquez, José L. Risco, Silvia Corrochano, Jose L. Ayala

**Affiliations:** Department of Computer Architecture and Automation, UCM, Madrid, Spain; Neurological Disorders Group, Hospital Clinico San Carlos, IdISSC, Madrid, Spain

**Keywords:** Amyotrophic lateral sclerosis, lifestyle questionnaires, large language models, natural language processing, longitudinal change, machine learning

## Abstract

**Background:** Mixed-format lifestyle questionnaires contain both structured variables and free-text responses, but it remains unclear whether language-derived variables provide incremental predictive value beyond structured data, and under which representational condition. It was investigated whether variables derived from patient-reported free text improve ALS-versus-control classification beyond structured questionnaire data, and whether their value depends on how temporal information is represented.

**Methods:** A leakage-free machine-learning pipeline was developed to classify ALS versus controls from questionnaire-derived data, including a schema-guided LLM-based text-to-table extraction and a compact longitudinal encoding strategy. Three feature configurations were compared: Pool1, containing structured baseline variables only; Pool2, adding compact summaries derived from first-time-point (T1) free-text responses; and Pool3, further incorporating compact descriptors of change between T1 and T2. Logistic Regression, linear Support Vector Classification, and Random Forest were evaluated using repeated stratified holdout (10 seeds) and repeated stratified 5-fold cross-validation. Final ablation analyses were performed to isolate the contribution of the compact text block and the compact temporal block.

**Results:** After leakage correction, performance estimates became more conservative, indicating that previous results had been optimistic. In the final configuration, Pool3 achieved the best performance, with Random Forest reaching a holdout accuracy of 0.673, F1-weighted score of 0.666, and Matthews correlation coefficient of 0.323; cross-validated F1-weighted score and Matthews correlation coefficient were 0.654 and 0.312, respectively. Pool2 did not show a robust improvement over Pool1. Ablation analysis showed that removing the compact temporal block markedly reduced Pool3 performance, whereas removing the compact text block had little overall effect. These findings indicate that the primary value of language-based processing in small clinical cohorts lies not in static feature enrichment, but in enabling compact representations of longitudinal change.

**Conclusions:** In this setting, the main predictive gain did not arise from static text-derived variables alone, but from representing questionnaire information as compact longitudinal change descriptors. These findings suggest that, in small clinical cohorts, the value of language-based processing may lie more in summarizing trajectories than in expanding static feature spaces.

## 1. Introduction

Amyotrophic lateral sclerosis (ALS) is a progressive and fatal neurodegenerative disorder characterized by the degeneration of upper and lower motor neurons, leading to progressive weakness, functional decline, and premature death [1,2]. Although ALS has traditionally been approached as a motor neuron disease, it is increasingly understood as a heterogeneous multisystem disorder that overlaps clinically and biologically with frontotemporal degeneration [2,3]. In parallel with the study of genetic mechanisms, considerable interest has focused on non-genetic contributors, including lifestyle and environmental exposures, as potential sources of phenotypic heterogeneity and disease-related signal [4]. This makes ALS a particularly suitable setting in which to study whether questionnaire-derived lifestyle information can contribute to phenotype classification.

In clinical and epidemiological research, lifestyle and exposure histories are often collected through mixed-format questionnaires that combine structured items with open-ended narrative responses. These patient-generated data can capture contextual information on habits, environment, and lived experience that is difficult to represent through fixed-choice variables alone [5]. However, once collected, much of this information remains semi-structured or unstructured, which limits its systematic integration into tabular analytical workflows and downstream machine learning models [5,6]. As a result, a potentially informative portion of questionnaire data is often underused unless it is manually coded, which is slow, labor-intensive, and difficult to standardize.

Natural language processing (NLP) and, more recently, large language models (LLMs) offer a possible solution to this problem by transforming free text into structured variables. In clinical informatics, a growing body of work has shown that NLP and LLM-based systems can successfully extract socially and behaviorally relevant information from unstructured clinical notes and convert it into database-ready representations [6-9]. Yet most of this evidence comes from provider-authored electronic health record narratives rather than from patient-reported lifestyle questionnaires. This distinction matters because patient-reported text differs in vocabulary, completeness, and semantic granularity, and because mixed-format questionnaires already contain a structured backbone against which the added value of text-derived variables must be judged.

A second limitation in the existing literature is that many studies emphasize extraction performance or documentation completeness, while relatively few evaluate whether text-derived variables improve downstream predictive modeling once existing structured data are already available [6-10]. This question is especially relevant in small clinical cohorts, where adding more variables may increase complexity without necessarily improving predictive signal. At the same time, recent methodological work on longitudinal clinical data suggests that predictive benefit may depend not only on the presence of additional information, but on how that information is represented over time [11]. In this context, language-based processing may be most useful not as a generator of richer static features but as a tool to summarize behavioral or exposure trajectories in a compact and model-compatible way.

The present study addresses these gaps in the context of ALS versus control classification from mixed-format lifestyle questionnaires. A leakage-free pipeline was developed to convert free-text questionnaire responses into structured variables and compare their predictive contribution across three increasingly enriched data configurations: structured baseline variables only (Pool1), baseline variables plus information derived from first-time-point narratives (Pool2), and baseline variables plus compact descriptors of change between the first and second time points (Pool3). The present study was designed not only to evaluate whether language-derived information improves prediction, but to test the hypothesis that its predictive value depends on how it is represented; specifically, whether it is incorporated as static feature enrichment or as compact summaries of longitudinal change. We hypothesize that compact longitudinal representations will provide greater predictive value than static text-derived features in small clinical cohorts.

## 2. State of the Art

Machine learning (ML) applications in amyotrophic lateral sclerosis (ALS) have grown steadily in recent years, but the field has been dominated by studies based on biomedical signals, neuroimaging, molecular biomarkers, and disease progression modeling rather than by patient-reported lifestyle questionnaires. A systematic review of ML in ALS identified diagnosis, communication support, and survival prediction as the main application areas, with most studies relying on signal-based or instrument-derived data [12]. More recent studies have extended this line of work to progression prediction from longitudinal clinical features [13] and to objective estimation of disease severity from voice and accelerometer recordings [14]. Taken together, this literature shows that ML can capture clinically relevant signal in ALS, but it has largely focused on structured or sensor-derived data rather than on mixed-format questionnaire data containing both closed and open-ended responses.

When questionnaire-derived variables have been used in ALS-related ML, the emphasis has generally been placed on functional status, caregiver burden, or quality of life rather than on lifestyle and exposure profiling. For example, ML models have been applied to predict caregiver burden and caregiver quality of life using demographic, socioeconomic, psychological, and clinical questionnaire variables collected in ALS cohorts [15,16]. Likewise, longitudinal ML approaches have been used to estimate ALS severity from repeated symptom-related and functional measures [14]. These studies are relevant because they demonstrate that questionnaire-based and longitudinal information can support predictive modeling in ALS. However, they do not address the specific question considered here: whether patient-reported lifestyle questionnaires, especially those containing free-text answers, provide added predictive value for distinguishing ALS from controls beyond what is already captured by structured variables.

Outside ALS, a large body of clinical NLP research has shown that unstructured text can be transformed into structured variables describing social, behavioral, and contextual health information. Systematic reviews have documented the extensive use of NLP for extracting social determinants of health (SDoH) from clinical narratives, especially smoking, alcohol use, substance use, housing, employment, and social support [6]. More recently, large language models have shown strong performance for extracting such information from free-text clinical notes, including across institutions and note types [7-10,17]. However, most of this literature has three features that distinguish it from the present setting. First, it predominantly uses provider-authored electronic health record notes rather than patient-generated questionnaire narratives. Second, it often evaluates extraction quality itself rather than the downstream predictive value of the derived variables once structured covariates are already available. Third, it is usually framed around documentation recovery or SDoH surveillance, not around the comparative value of questionnaire blocks in a disease-classification pipeline.

A second relevant body of literature concerns predictive modeling with longitudinal clinical data. Methodological reviews have shown that artificial intelligence models increasingly exploit longitudinal EHR information, but they have also highlighted substantial heterogeneity in temporal encoding, limited methodological transparency, frequent reliance on a single train-test split, and generally weak reporting practices [11]. This is important because the predictive usefulness of repeated measures depends not only on the availability of follow-up data, but also on how temporal information is represented. In other words, simply appending a second time point may increase dimensionality without improving signal, whereas compact representations of change may better capture clinically meaningful trajectories. This point is particularly relevant in small datasets, where the balance between information richness and model complexity is especially delicate.

Against this background, the specific gap addressed by the present work lies at the intersection of four areas: ML in ALS, questionnaire-based phenotyping, free-text information extraction, and longitudinal representation. The literature reviewed above suggests that ML in ALS is already established, that NLP and LLMs are effective for structuring unstructured health-related text, and that longitudinal representations can provide added predictive value when encoded appropriately. However, evidence remains limited on whether free-text responses from patient lifestyle questionnaires improve ALS-versus-control classification once structured questionnaire data are already available, and even more limited on whether the main value of language-based processing lies in static feature enrichment or in compact summaries of longitudinal change. This study was designed specifically to address that gap by comparing structured baseline data (Pool1), baseline plus text-derived first-time-point variables (Pool2), and baseline plus compact longitudinal descriptors between T1 and T2 (Pool3).

## 3. Methods

### 3.1. Study design, ethics, data source, and cohort definition

A retrospective observational study was conducted using questionnaire-derived lifestyle and clinical data collected at Hospital Clínico San Carlos. The study was approved by the local ethical committee at the Hospital Clinico San Carlos (19-524-E), and all patients and healthy controls read and signed the informed consent to participate in the study. The source dataset was delivered as a Microsoft Excel workbook containing a data dictionary and a consolidated sheet with baseline variables, time-point-specific variables, and free-text interview fields. The original workbook contained multiple diagnostic groups. For the present binary-classification study, the target label was restricted to Control and ALS. Records labelled as ALS-FTD were relabelled as ALS in order to preserve sample size and remain aligned with the primary classification objective, whereas records corresponding to FTD were excluded from the main analyses. After label harmonization and cohort filtering, the final analytic sample comprised 103 participants (60 controls and 43 ALS cases).

### 3.2. Questionnaire structure, temporal framing, and pool definitions

The questionnaire combined structured items and open-ended narrative responses. Information was collected in two temporal frames: time point 1 (T1), corresponding to earlier life, and time point 2 (T2), corresponding to later/current life. The temporal interpretation depended on age: for participants younger than 65 years, T1 referred to experiences up to 35 years of age and T2 to the period after 35 years; for participants aged 65 years or older, T1 referred to the period before 65 years and T2 to the post-retirement period. The questionnaire covered baseline demographic and medical-history variables, structured lifestyle variables at T1 and/or T2 (e.g., housing, education, socioeconomic context, work, physical activity, diet, sleep, and substance use), and several free-text fields describing work history, sport habits, diet, social life, substance use, and general interview content.

Three experimental feature configurations were defined. Pool1 contained structured baseline variables only. Pool2 extended Pool1 with structured T1 variables and compact summaries derived from T1 free-text content. Pool3 extended Pool2 with structured T2 information and compact descriptors of change between T1 and T2. All three pools were retained as comparative interpretation: Pool1 as the structured baseline, Pool2 as the static text-enriched configuration, and Pool3 as the longitudinally enriched configuration. The age-dependent definition of T1 and T2 is illustrated in **Figure 1**.

**Figure 1.**
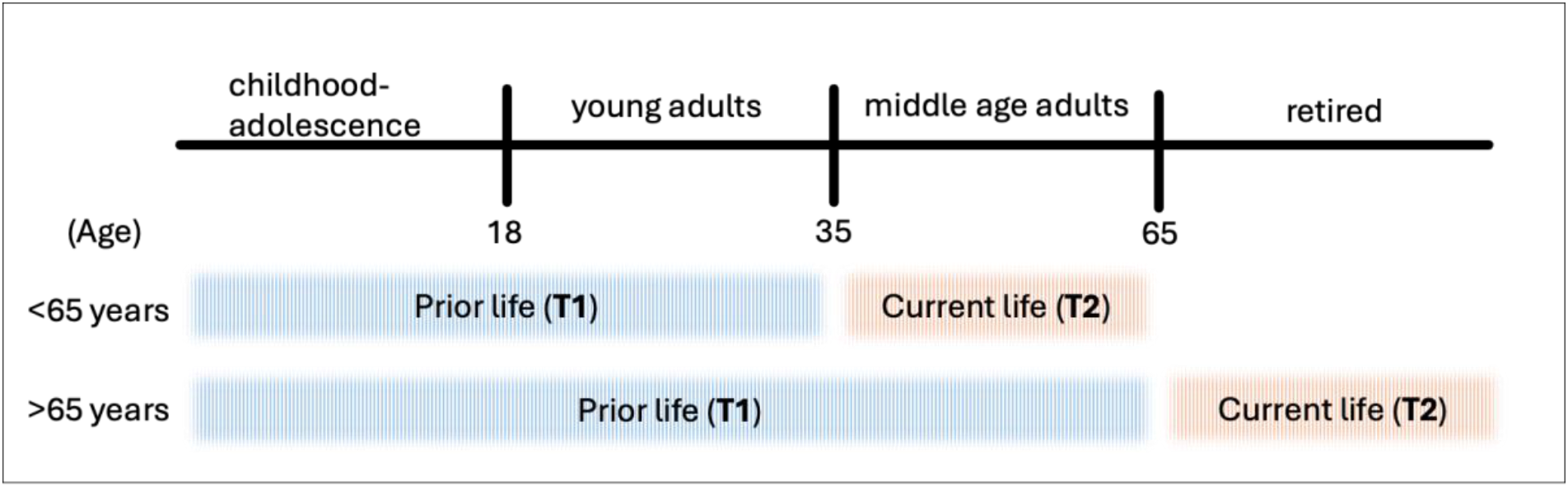
Temporal framing of questionnaire information used to define T1 and T2 according to participant age. For participants younger than 65 years, T1 corresponds to experiences up to 35 years of age and T2 to the period after 35 years. For participants aged 65 years or older, T1 corresponds to the period before age 65 and T2 to the post-retirement period.

The overall study workflow, from source questionnaire data and cohort definition to LLM-based feature extraction, compact feature construction, leakage-free preprocessing, model training, and evaluation, is summarized in **Figure 2**.

**Figure 2.**
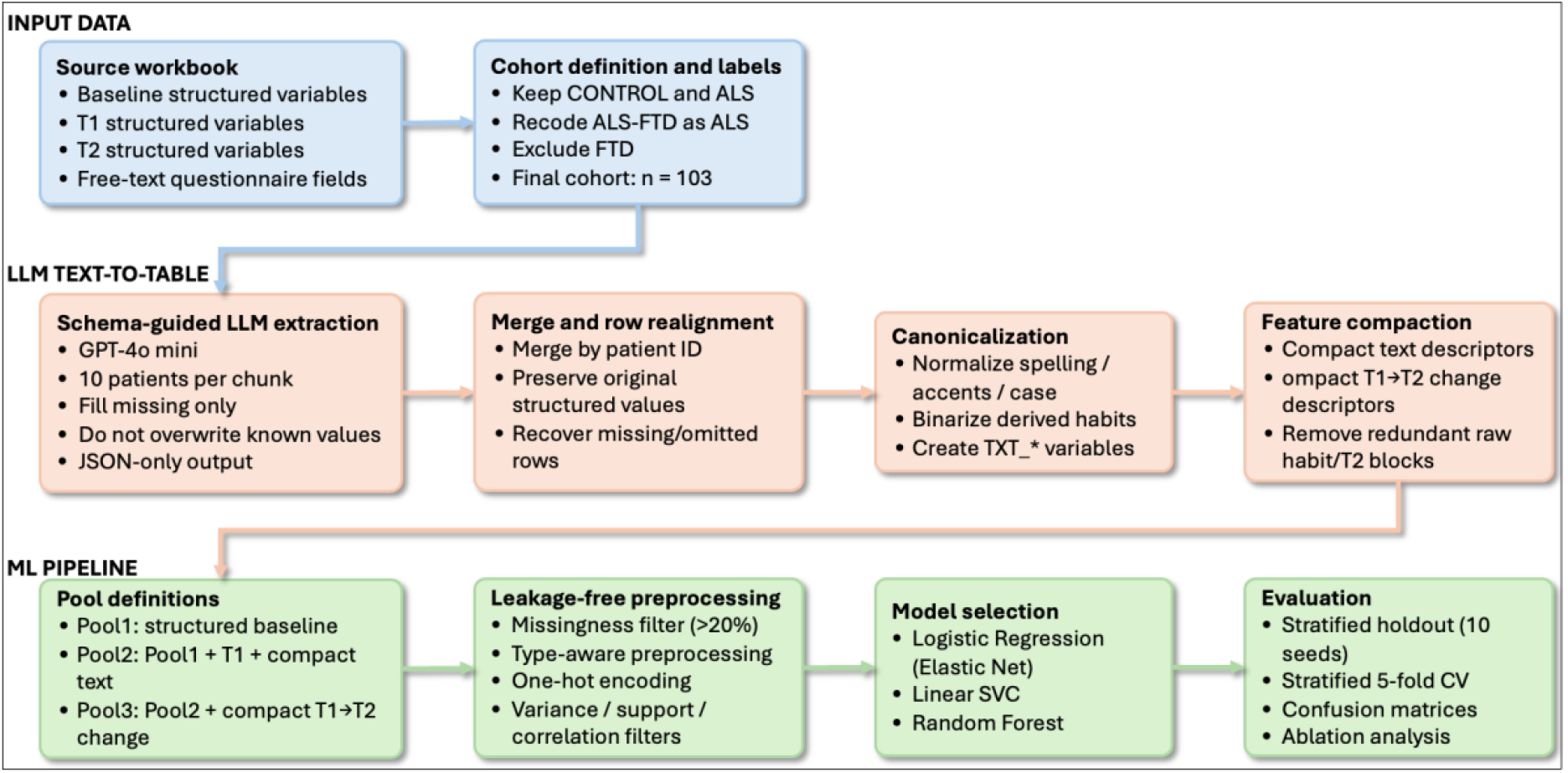
Final leakage-free study workflow. Overview of the full analytical pipeline, including source questionnaire data, cohort definition, schema-guided LLM-based text-to-table extraction, row realignment, canonicalization of text-derived variables, construction of compact feature blocks, pool definition, leakage-free preprocessing, model selection, and repeated evaluation by stratified holdout and stratified cross-validation.

### 3.3. Initial variable harmonization

Before any language-model processing, several variables were harmonized to reduce redundancy and align feature types. Age was discretized into three bins (≤35, 36-65, >65 years), consistent with the temporal design of the questionnaire. BMI at T1 and T2 was transformed into categorical variables using standard adult BMI cutoffs (underweight, normal weight, overweight, obesity) based on World Health Organization thresholds [18]. The original continuous age and BMI columns were then removed.

Additional harmonization steps were applied to avoid duplicated semantics across columns. For example, sex-specific formulations of body-shape variables were coalesced into unified variables (BODYSHAPE1 and BODYSHAPE2), and minor column-name inconsistencies (e.g., WEIGHTTYPE_2) were normalized. These operations were performed before model evaluation because they represent semantic harmonization rather than statistical learning from the outcome.

### 3.4. LLM-based text-to-table enrichment

#### 3.4.1. Task formulation

Free-text questionnaire responses were transformed into structured variables through a schema-guided generative information extraction workflow, conceptually aligned with text-to-table extraction [19,20]. Rather than generating free-form summaries, the model was instructed to return JSON objects containing patient-level updates to a predefined structured schema. The goal of this stage was twofold: (i) to complete missing structured lifestyle values only when clearly supported by the text, and (ii) to identify recurrent lifestyle habits that were repeatedly expressed in the narratives but not explicitly represented in the original tabular schema. The schema-guided extraction workflow, from free-text questionnaire responses to structured JSON outputs and compact text descriptors, is summarized in **Figure 3**.

**Figure 3.**
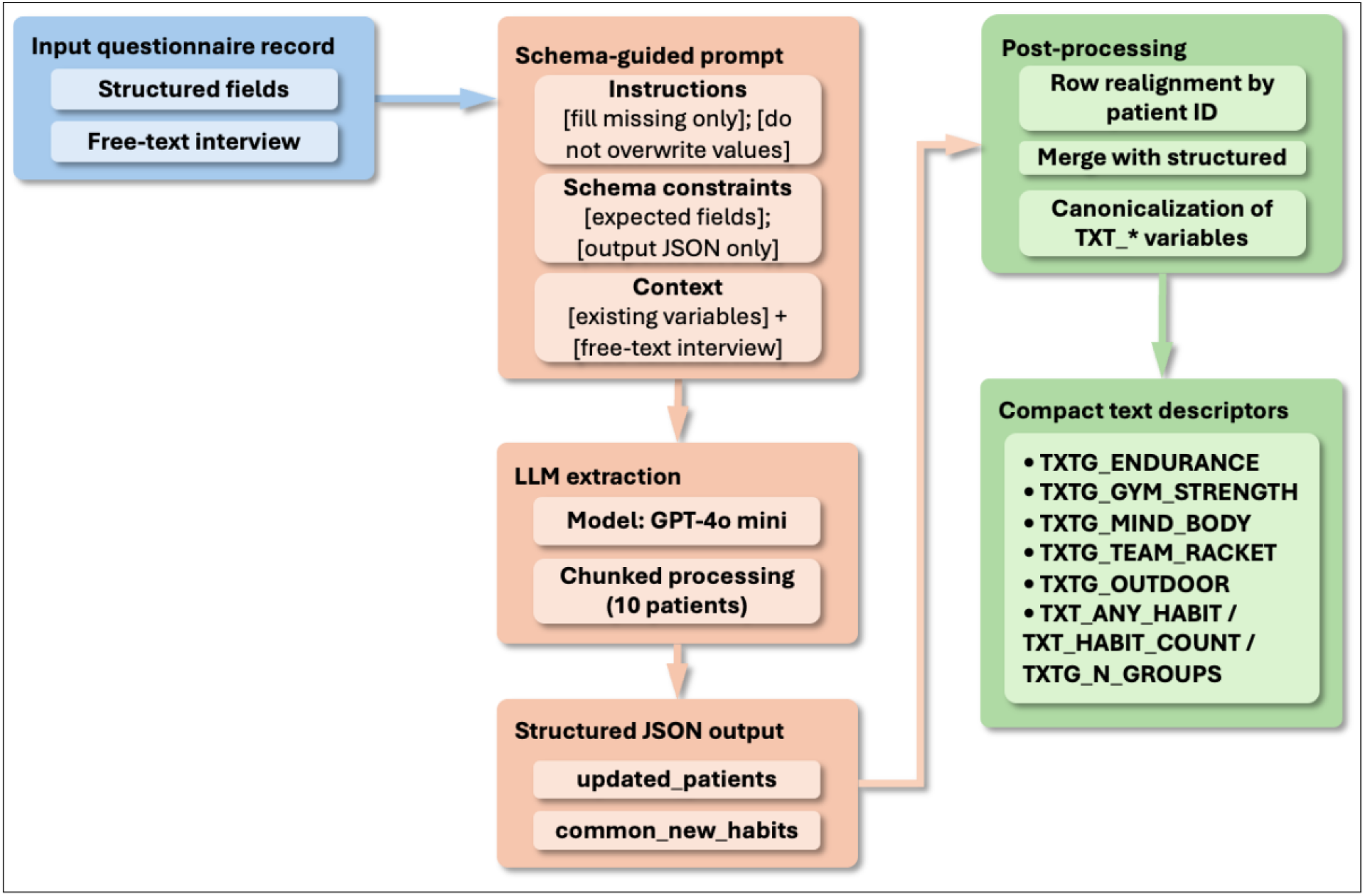
Schema-guided LLM extraction from free-text questionnaire responses and post-processing into compact text descriptors. Structured questionnaire fields and free-text interview responses were combined with a constrained prompt to generate JSON-formatted updates. The outputs were then realigned by patient identifier, canonicalized, and aggregated into compact text-summary variables for downstream modeling.

**Figure 4.** Construction of compact temporal descriptors and leakage-free model development. Panel A illustrates the generation of compact temporal descriptors from paired T1 and T2 variables, including selected DELTA_* variables and domain-level T12_* summaries. Panel B illustrates the leakage-free preprocessing and evaluation strategy, in which missingness filtering, preprocessing, encoding, unsupervised filtering, hyperparameter tuning, and model fitting were performed exclusively within the training partitions before application to validation or test data.

#### 3.4.2. Prompt design and API implementation

The extraction prompt defined the model as a conservative “data assistant” and enforced strict output rules: only complete missing structured values when explicit textual evidence existed, never overwrite values already present in the structured data, preserve previously known candidate habits, and return JSON-only output. The prompt, therefore, combined extraction instructions, schema context, and format constraints.

Model calls were implemented through the OpenAI Chat Completions API using GPT-4o mini, with temperature = 0.2 to reduce response variability and encourage conservative completion behavior [21,22]. To remain within context-window constraints and to stabilize extraction, the dataset was processed in chunks of 10 patients per request. No manual post-hoc correction of LLM outputs was performed beyond deterministic canonicalization rules.

The extraction process was not formally evaluated against a human-annotated gold standard, as the objective of this study was downstream predictive utility rather than extraction accuracy.

#### 3.4.3. Chunked processing, row realignment, and canonicalization

Each API call returned two elements: an updated patient list and a list of potentially new common habits. To ensure consistency across chunks, newly proposed habits were appended to a mutable list of candidate habit columns and provided back to the model in subsequent requests. Because language-model outputs may omit records or return incomplete chunks, a row-realignment step was introduced to merge generated outputs back to the original patient table by patient identifier and to preserve all pre-existing structured values. Missing returned records were therefore reconstructed from the original structured input rather than discarded.

After chunk-wise merging, free-text-derived habit variables were canonicalized. This involved normalization of accents, case, and spelling variants, conversion to a unified TXT_* naming convention, and binarization of presence/absence values. Habit-related missing values were treated as absence when conceptually appropriate, thereby avoiding the propagation of empty fields for purely binary text-derived indicators.

### 3.5. Compact representation of text-derived information

The intermediate text-derived variables were then summarized into a reduced set of compact descriptors. Instead of retaining each individual free-text habit as an independent feature, conceptually related habits were grouped into five broader categories: TXTG_ENDURANCE (e.g., walking, running, cycling, swimming, athletics, hiking); TXTG_GYM_STRENGTH (e.g., gym, cross-fit, boxing, gymnastics); TXTG_MIND_BODY (e.g., yoga, pilates, tai chi, dance); TXTG_TEAM_RACKET (e.g., football, basketball, handball, volleyball, tennis, padel) and TXTG_OUTDOOR (e.g., walking, running, cycling, hiking, golf).

In addition, three summary variables were created: TXT_ANY_HABIT, indicating whether any free-text habit was present; TXT_HABIT_COUNT, indicating the total number of individual text-derived habits detected; and TXTG_N_GROUPS, indicating the number of compact habit groups present. After this step, the individual TXT_* habit columns were removed from the final compact representation. This reformulation was designed to reduce fragmentation of the text-derived signal and improve its compatibility with downstream machine learning.

This design explicitly prioritizes dimensionality reduction and semantic aggregation over feature granularity in order to mitigate overfitting in small-sample settings.

### 3.6. Compact longitudinal representation of change between T1 and T2

In Pool3, T2 was not introduced as a second raw feature block. Instead, the final pipeline summarized longitudinal evolution through compact descriptors of change between T1 and T2. First, paired T1/T2 variables were automatically identified through variable-name matching. Second, for each paired variable, the pipeline computed whether the value changed between time points. Third, these patient-level changes were summarized through domain-specific descriptors.

Two types of temporal features were created. The first type consisted of global and domain-level change counts/proportions, such as the total number of observed T1/T2 pairs, the total number of changed pairs, and the number or proportion of changes within predefined domains (socioeconomic, activity, body-related, substance/sleep, psychosocial, and diet). The second type consisted of selected delta variables for specific interpretable measures, including body-related and behavioral variables such as weight, height, sport frequency, sleep hours, concern, personality, socialization, dietary profile, sugary drinks, sweets, fast food, number of meals, food quantity, and satiety. Additional global descriptors summarizing the magnitude of change (mean absolute delta and maximum absolute delta) were also generated. After these compact longitudinal descriptors had been created, the raw T2 columns were removed from the final Pool3 representation to reduce redundancy and effective dimensionality.

Importantly, this approach reframes longitudinal information as change rather than state, reducing redundancy and emphasizing the trajectory-level signal.

### 3.7. Leakage-free preprocessing and feature representation

All data-dependent preprocessing steps were embedded within the training partitions of the validation workflow in order to prevent information leakage. First, textual entries were sanitized, normalized, and coerced into numeric-compatible formats whenever appropriate. Within each training partition, variables with more than 20% missingness were removed. The remaining predictors were then processed according to variable type using a custom preprocessor that separated binary, categorical, and numeric features. Binary and categorical variables were imputed with the most frequent value, whereas numeric variables were imputed using the median and subsequently standardized. Categorical predictors were then one-hot encoded, with infrequent levels grouped using min_frequency = 3 in order to avoid the generation of isolated dummy variables. After encoding, predictors with zero variance were discarded, followed by the removal of features with fewer than three non-zero observations and the filtering of highly correlated predictors using an absolute pairwise correlation threshold of 0.90. Variables belonging to the compact temporal block (DELTA_*, T12_*) and count-based summary variables (TXT_HABIT_COUNT, TXTG_N_GROUPS) were explicitly treated as numeric variables to avoid inappropriate one-hot expansion.

### 3.8. Classification models and hyperparameter tuning

Three supervised classifiers were evaluated: Logistic Regression, linear Support Vector Classification, and Random Forest. Logistic Regression was implemented with the saga solver and Elastic Net regularization, which allows a continuum between L1-like sparsity and mixed L1/L2 penalization [23]. Linear SVC was chosen as a strong baseline for high-dimensional classification settings [24]. Random Forest was included as a non-linear ensemble model capable of capturing variable interactions [25].

Hyperparameters were tuned within each training partition using an inner 3-fold stratified cross-validation procedure and macro-F1 as the tuning metric. For Logistic Regression, the grid included C = {0.001, 0.01, 0.1, 1, 10}, l1_ratio = {0.5, 0.8, 1.0}, and class_weight ∈ {None, balanced}. For linear SVC, the grid included C = {0.001, 0.01, 0.1, 1, 10} and class_weight ∈ {None, balanced}. For Random Forest, a randomized search was performed over n_estimators ∈ {100, 200, 300}, max_depth ∈ {None, 3, 5, 8}, min_samples_split ∈ {2, 5, 10}, min_samples_leaf ∈ {1, 2, 4, 6}, max_features ∈ {sqrt, 0.2, 0.5}, and class_weight ∈ {None, balanced, balanced_subsample}.

### 3.9. Evaluation strategy and performance metrics

Model performance was assessed through two complementary repeated stratified evaluation schemes. First, a repeated stratified 5-fold cross-validation was run using 10 fixed random seeds (42-51), yielding 50 outer folds per model and pool. Second, a repeated stratified holdout procedure was performed using the same 10 seeds and a 75/25 train-test split, with hyperparameter tuning repeated inside each training split. This design provided both a repeated test-set perspective and a repeated cross-validation perspective, while preserving strict train-test separation in all data-dependent steps.

The primary reported metrics were accuracy, weighted F1, F1 for the ALS class, ALS recall, balanced accuracy, and Matthews correlation coefficient (MCC). For interpretability, confusion matrices were aggregated across splits/folds, and false positives/false negatives were explicitly monitored. The target labels were encoded as binary values after merging ALS-FTD into the ALS class.

### 3.10. Ablation analysis

To isolate the contribution of the text-derived and longitudinal blocks, a final ablation analysis was performed on the best-performing configurations. In Pool2, the best model (Logistic Regression) was evaluated in two scenarios: the full model and the model with the compact text block removed. In Pool3, the best model (Random Forest) was evaluated in four scenarios: full model, without the compact text block, without the compact temporal block, and without either block. This analysis was designed to determine whether the observed gain in Pool3 was primarily attributable to the static text summaries, the temporal summaries, or both.

### 3.11. Reproducibility and implementation

All preprocessing, modeling, and evaluation steps were implemented in Python using scikit-learn pipelines and model-selection utilities [26]. The leakage-free design ensured that missingness filtering, preprocessing, dimensionality reduction, and hyperparameter selection were re-estimated independently within each training partition. Intermediate LLM-generated tables were exported for traceability, but only the fold-specific training data were used to fit preprocessing components and models during evaluation.

## 4. Results

A total of 103 participants were included in the binary classification experiments (60 controls and 43 ALS cases). Three data configurations were evaluated throughout the study: Pool1, containing structured baseline variables only; Pool2, adding variables derived from first-time-point free-text responses; and Pool3, further incorporating second-time-point information. Model performance was assessed using repeated stratified holdout evaluation across 10 random seeds and repeated stratified 5-fold cross-validation across the same seeds. Three successive optimization stages were performed, followed by a final ablation analysis to isolate the contribution of the compact text block and the compact temporal block. Pool3 was retained as the final pipeline, whereas Pool1 and Pool2 were kept as essential comparative conditions to determine which components of the questionnaires were actually contributing to predictive performance.

### 4.1. Stage-wise optimization of the predictive pipeline

The first optimization stage focused on removing information leakage by moving imputation, encoding, dimensionality reduction, and model fitting entirely inside the validation workflow. This resulted in a marked decrease in performance estimates, indicating that previous analyses had been overly optimistic. A second stage then addressed the consistency of the text-derived variables and corrected row-alignment issues between LLM-derived outputs and patient-level records. These additional cleaning steps produced only minor quantitative changes and did not alter the overall interpretation. The third stage introduced a compact representation of free-text content and, in Pool3, replaced raw T2 variables with compact descriptors of change between T1 and T2. This final reformulation produced the clearest performance gain, particularly in Pool3.

As shown in both **Table 1A** and **Table 1B**, after leakage correction, neither Pool2 nor Pool3 consistently outperformed Pool1. Additional cleaning of text-derived variables also failed to materially modify this pattern. In contrast, the final compact reformulation substantially improved Pool3, whose best holdout F1-weighted score increased to 0.666 and whose best cross-validated F1-weighted score increased to 0.654. By comparison, Pool2 remained comparatively stable across iterations, supporting the interpretation that first-time-point free-text information alone did not provide a robust improvement over the structured baseline.

**Table 1.**
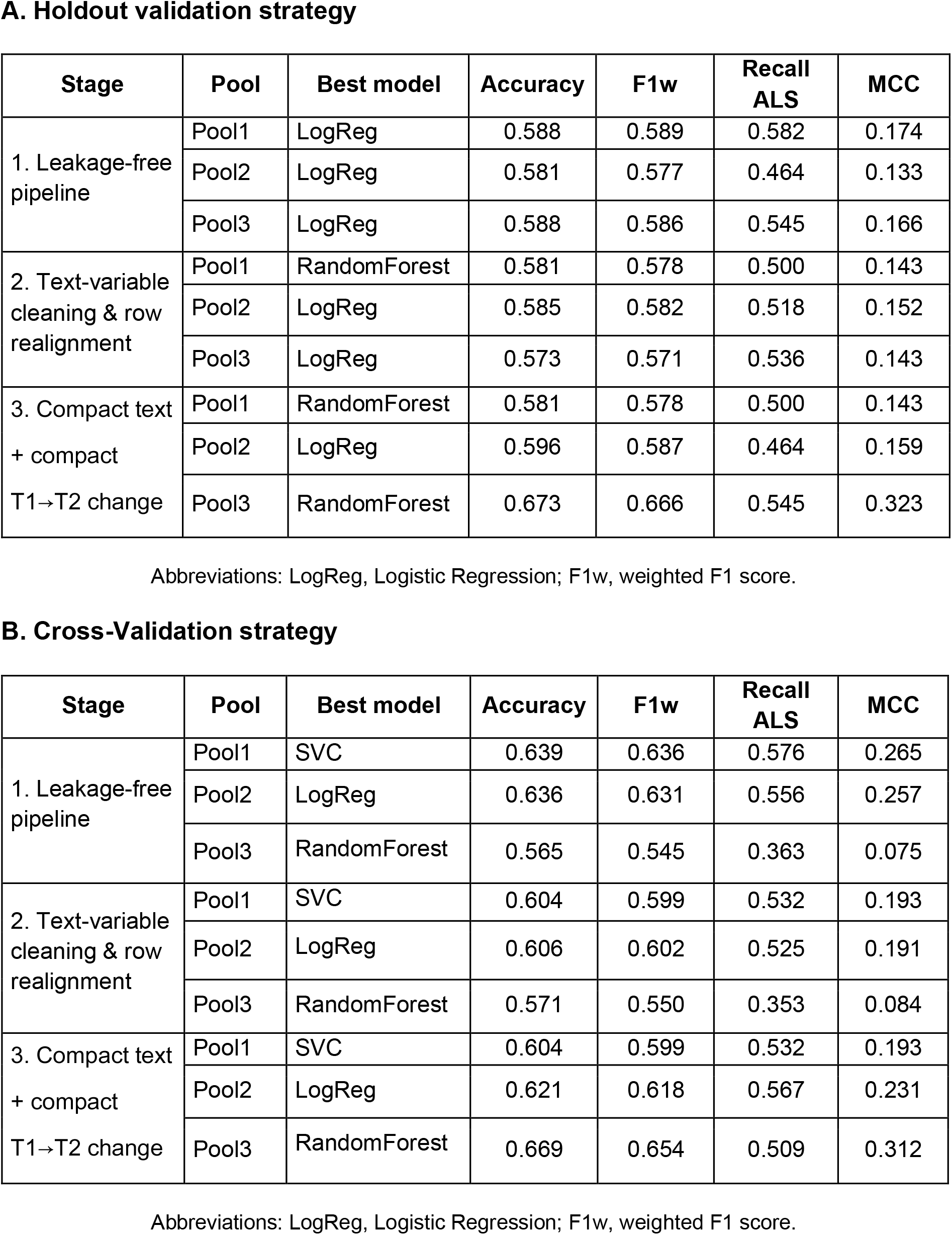
Evolution of the best-performing configuration of each pool across the three optimization stages. **(A)** Holdout metrics correspond to the mean performance across 10 stratified splits. **(B)** Cross-validation metrics correspond to repeated stratified 5-fold cross-validation across the same seeds.

A. Holdout validation strategy
| Stage | Pool | Best model | Accuracy | F1w | Recall ALS | MCC |
| --- | --- | --- | --- | --- | --- | --- |
| 1. Leakage-free pipeline | Pool1 | LogReg | 0.588 | 0.589 | 0.582 | 0.174 |
|  | Pool2 | LogReg | 0.581 | 0.577 | 0.464 | 0.133 |
|  | Pool3 | LogReg | 0.588 | 0.586 | 0.545 | 0.166 |
| 2. Text-variable cleaning & row realignment | Pool1 | RandomForest | 0.581 | 0.578 | 0.500 | 0.143 |
|  | Pool2 | LogReg | 0.585 | 0.582 | 0.518 | 0.152 |
|  | Pool3 | LogReg | 0.573 | 0.571 | 0.536 | 0.143 |
| 3. Compact text + compact T1→T2 change | Pool1 | RandomForest | 0.581 | 0.578 | 0.500 | 0.143 |
|  | Pool2 | LogReg | 0.596 | 0.587 | 0.464 | 0.159 |
|  | Pool3 | RandomForest | 0.673 | 0.666 | 0.545 | 0.323 |

Abbreviations: LogReg, Logistic Regression; F1w, weighted F1 score.

B. Cross-Validation strategy
| Stage | Pool | Best model | Accuracy | F1w | Recall ALS | MCC |
| --- | --- | --- | --- | --- | --- | --- |
| 1. Leakage-free pipeline | Pool1 | SVC | 0.639 | 0.636 | 0.576 | 0.265 |
|  | Pool2 | LogReg | 0.636 | 0.631 | 0.556 | 0.257 |
|  | Pool3 | RandomForest | 0.565 | 0.545 | 0.363 | 0.075 |
| 2. Text-variable cleaning & row realignment | Pool1 | SVC | 0.604 | 0.599 | 0.532 | 0.193 |
|  | Pool2 | LogReg | 0.606 | 0.602 | 0.525 | 0.191 |
|  | Pool3 | RandomForest | 0.571 | 0.550 | 0.353 | 0.084 |
| 3. Compact text + compact T1→T2 change | Pool1 | SVC | 0.604 | 0.599 | 0.532 | 0.193 |
|  | Pool2 | LogReg | 0.621 | 0.618 | 0.567 | 0.231 |
|  | Pool3 | RandomForest | 0.669 | 0.654 | 0.509 | 0.312 |

Abbreviations: LogReg, Logistic Regression; F1w, weighted F1 score.

These results suggest that neither additional variables nor improved preprocessing alone were sufficient to enhance performance, and that representational reformulation was the key driver of improvement.

### 4.2. Final compact representation identified Pool3 as the best-performing configuration

The third optimization stage was retained as the final comparative configuration because it provided the most informative representation of the questionnaires. In this stage, Pool1 remained the structured baseline, Pool2 incorporated compact summaries of free-text information from T1, and Pool3 additionally included compact longitudinal descriptors of change between T1 and T2. The detailed holdout performance of all three models across the three pools is shown in **Table 2**.

**Table 2.** Holdout performance of the final compact representation for each model and pool. Values correspond to the mean across 10 stratified splits. Pool1 contains structured baseline variables only; Pool2 adds compact summaries derived from T1 free-text responses; and Pool3 further incorporates compact descriptors of change between T1 and T2. Abbreviations: LogReg, Logistic Regression; F1w, weighted F1 score. Values correspond to the mean across 10 stratified splits.

| Pool | Model | Accuracy | F1w | F1 ALS | Recall ALS | Balanced Accuracy | MCC |
| --- | --- | --- | --- | --- | --- | --- | --- |
| Pool1 | LogReg | 0.573 | 0.570 | 0.484 | 0.482 | 0.561 | 0.122 |
|  | SVC | 0.546 | 0.545 | 0.463 | 0.464 | 0.535 | 0.071 |
|  | RF | 0.581 | 0.578 | 0.502 | 0.500 | 0.570 | 0.143 |
| Pool2 | LogReg | 0.596 | 0.587 | 0.483 | 0.464 | 0.578 | 0.159 |
|  | SVC | 0.577 | 0.569 | 0.465 | 0.455 | 0.561 | 0.120 |
|  | RF | 0.538 | 0.528 | 0.398 | 0.364 | 0.515 | 0.033 |
| Pool3 | LogReg | 0.573 | 0.568 | 0.499 | 0.509 | 0.565 | 0.136 |
|  | SVC | 0.596 | 0.592 | 0.518 | 0.527 | 0.587 | 0.175 |
|  | RF | 0.673 | 0.666 | 0.580 | 0.545 | 0.656 | 0.323 |

Within this final configuration, Pool3 clearly achieved the strongest performance, particularly with Random Forest, which reached an accuracy of 0.673 ± 0.091, an F1-weighted score of 0.666 ± 0.095, a balanced accuracy of 0.656 ± 0.096, a recall for ALS of 0.545 ± 0.155, and an MCC of 0.323 ± 0.200. In contrast, the best holdout F1-weighted values in Pool1 and Pool2 were 0.578 ± 0.091 and 0.587 ± 0.095, respectively. Thus, the final improvement was not explained by the mere addition of more variables, but rather by a more informative and more compact representation of longitudinal change.

The confusion matrices further supported this interpretation, as shown in **Figure 5**, which illustrates the reduction in both false positives and false negatives in the final Pool3 model and the deterioration observed when the temporal block is removed. Compared with the best structured baseline model in Pool1 (Random Forest; holdout confusion matrix: [[96, 54], [55, 55]]) and the full Pool2 model (Logistic Regression; holdout confusion matrix: [[104, 46], [59, 51]]), the final Pool3 model reduced both false positives and false negatives (Random Forest; holdout confusion matrix: [[115, 35], [50, 60]]).

**Figure 5.**
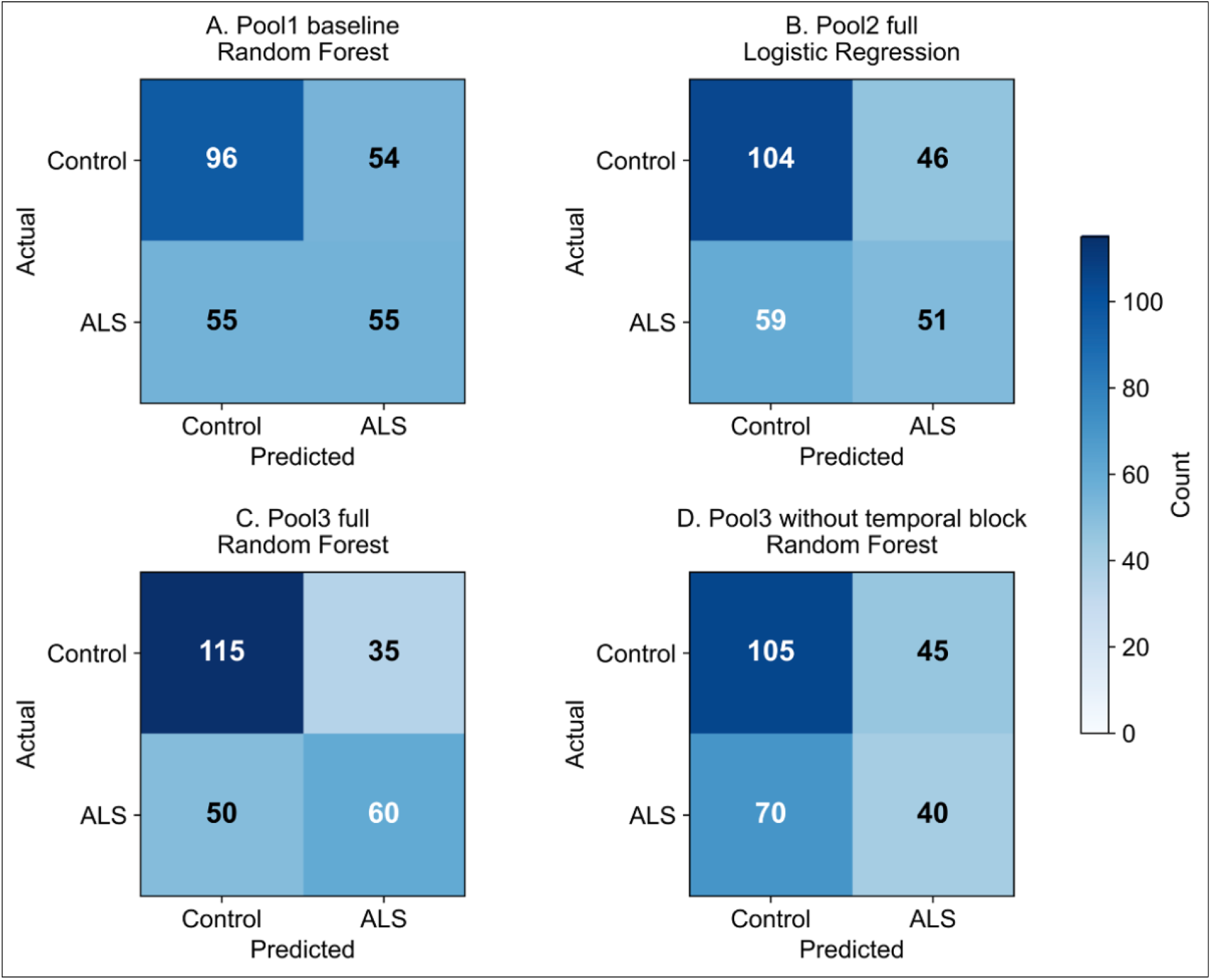
Holdout confusion matrices of the key final configurations. **(A)** Pool1 baseline (Random Forest), **(B)** Pool2 full model (Logistic Regression), **(C)** Pool3 full model (Random Forest), and **(D)** Pool3 without temporal block (Random Forest). Confusion matrices are shown as [[TN, FP], [FN, TP]].

This interpretation is also supported by the evolution of effective feature dimensionality across optimization stages, shown in **Figure 6**. While Pool1 remained low-dimensional and Pool2 showed only minor changes across stages, Pool3 exhibited a marked reduction in retained features in the final configuration. In particular, a diagnostic pre-compaction reference within the final pipeline yielded a mean of 212.9 retained features after filtering, whereas the final compact representation retained 148.2. This indicates that the improvement observed in Pool3 was associated not simply with the addition of T2 information, but with a more efficient and informative representation of longitudinal change.

**Figure 6.**
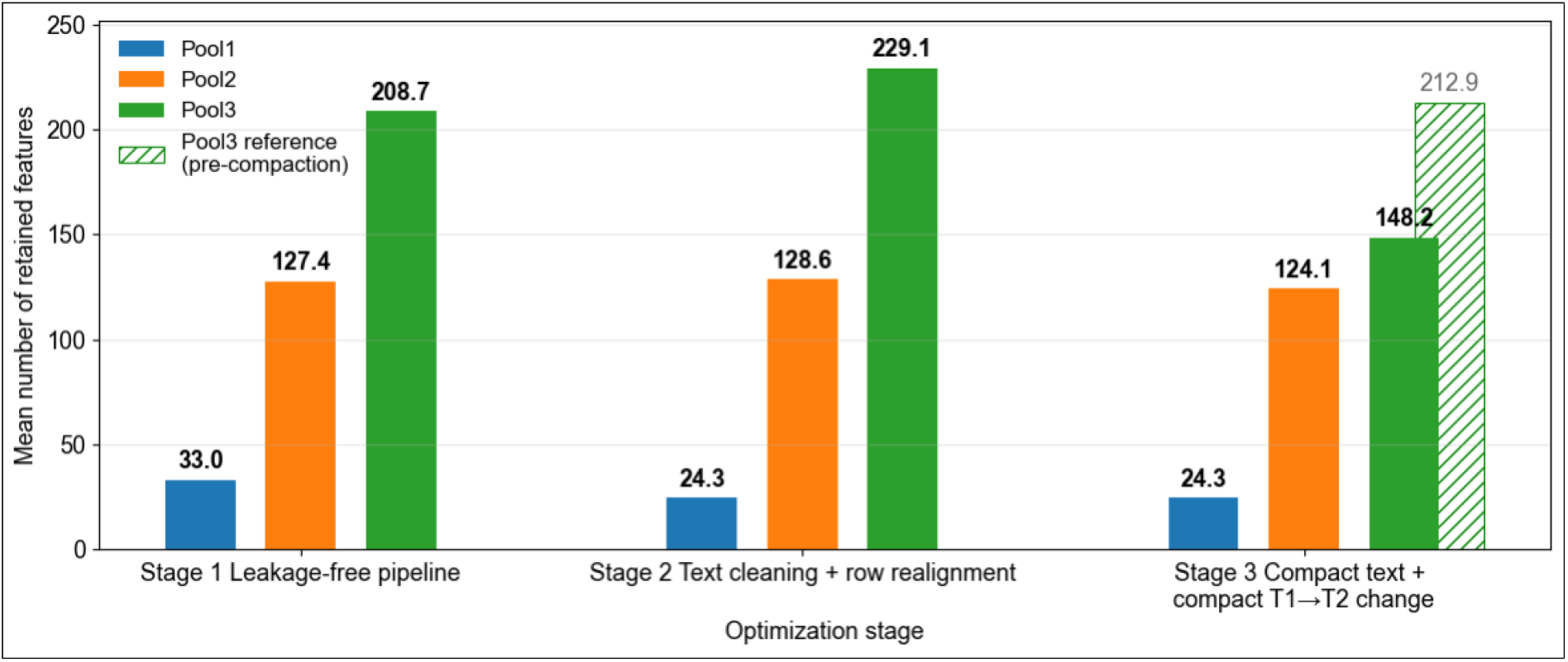
Retained feature dimensionality across optimization stages. Values correspond to the mean number of retained features across the repeated evaluation partitions after preprocessing and unsupervised filtering in each pool. The hatched bar in Stage 3 represents the real pre-compaction Pool3 reference measured within the final pipeline (n_after_correlation = 212.9), showing that compact reformulation of temporal information reduced the effective number of retained features to 148.2.

This indicates that performance gains were associated with improved information density rather than increased feature space.

### 4.3. Ablation analysis identified the compact temporal block as the main source of improvement

A final ablation analysis was performed to disentangle the contribution of the compact text block and the compact temporal block. In Pool2, the best model (Logistic Regression) was evaluated in two settings: full model and without the text block. In Pool3, the best model (Random Forest) was evaluated in four settings: full model, without text, without the temporal block, and without either block. The results are summarized in **Table 3**.

**Table 3.** Ablation analysis of the final models in Pool2 and Pool3. Holdout corresponds to the mean performance across 10 stratified splits; cross-validation corresponds to repeated stratified 5-fold cross-validation across the same seeds. Abbreviations: LogReg, Logistic Regression; F1w, weighted F1 score.

| Pool | Design | Holdout F1w | Holdout Recall ALS | Holdout MCC | CV F1w | CV Recall ALS | CV MCC |
| --- | --- | --- | --- | --- | --- | --- | --- |
| Pool 2 | Full model (LogReg) | 0.587 | 0.464 | 0.159 | 0.618 | 0.567 | 0.231 |
| Pool 2 | Without text block | 0.620 | 0.527 | 0.224 | 0.614 | 0.540 | 0.219 |
| Pool 3 | Full model (RF) | 0.666 | 0.545 | 0.323 | 0.654 | 0.509 | 0.312 |
| Pool 3 | Without text block | 0.683 | 0.536 | 0.368 | 0.654 | 0.532 | 0.310 |
| Pool 3 | Without temporal block | 0.542 | 0.364 | 0.066 | 0.558 | 0.399 | 0.098 |
| Pool 3 | Without text and temporal blocks | 0.594 | 0.436 | 0.191 | 0.573 | 0.406 | 0.137 |

Abbreviations: LogReg, Logistic Regression; F1w, weighted F1 score.

In Pool2, removing the text block improved holdout performance from an F1-weighted score of 0.587 to 0.620 and MCC from 0.159 to 0.224, but slightly reduced cross-validated performance from an F1-weighted score of 0.618 to 0.614 and MCC from 0.231 to 0.219. This pattern indicates that text-derived variables from T1 did not provide a robust and reproducible gain over the structured baseline. In Pool3, removing the text block had little overall effect and even slightly improved holdout performance, with F1-weighted increasing from 0.666 to 0.683 and MCC from 0.323 to 0.368, while cross-validated F1-weighted remained essentially unchanged (0.654 in both cases). In contrast, removing the temporal block caused a marked drop in all metrics: holdout F1-weighted decreased from 0.666 to 0.542 and MCC from 0.323 to 0.066, whereas cross-validated F1-weighted decreased from 0.654 to 0.558 and MCC from 0.312 to 0.098. These results indicate that the main contribution of the final Pool3 pipeline arises from the compact longitudinal representation rather than from the static text-derived variables themselves. Therefore, this pattern provides strong evidence that temporal encoding, rather than textual enrichment, is the dominant source of predictive signal in the final model.

The confusion matrices were consistent with this pattern. In Pool2, the full model yielded [[104, 46], [59, 51]] in holdout, whereas removing the text block yielded [[104, 46], [52, 58]], suggesting a reduction in false negatives but no change in false positives. In Pool3, the full model yielded [[115, 35], [50, 60]], while removing only the text block yielded [[121, 29], [51, 59]], indicating minimal practical differences. By contrast, removing the temporal block resulted in [[105, 45], [70, 40]], showing a substantial increase in false negatives and a clear deterioration in classification performance.

### 4.4. Summary of the comparative findings across pools

Overall, the comparative analysis supports three main findings. First, leakage correction substantially improved the methodological validity of the study by removing overly optimistic performance estimates. Second, the addition of text-derived variables from T1 alone did not provide a robust and reproducible improvement in classification performance. Third, the strongest gain emerged only after follow-up information was represented through compact longitudinal descriptors, with Pool3 consistently outperforming the other two pools in the final configuration. Accordingly, Pool3 was retained as the final pipeline for downstream interpretation, whereas Pool1 and Pool2 remained essential comparative conditions for identifying which components of the questionnaires were actually contributing to predictive performance.

## 5. Discussion

The present study yields three main findings. First, correcting information leakage substantially reduced the apparent performance of the models, but at the same time improved the methodological validity of the study. Second, variables derived from first-time-point free-text responses did not provide a robust and reproducible improvement over structured baseline variables alone. Third, the clearest gain emerged only after the follow-up information was reformulated as a compact representation of longitudinal change between T1 and T2, with Pool3 outperforming both Pool1 and Pool2 in the final configuration. The key contribution of this work is not the use of LLMs per se, but the demonstration that their utility in small clinical datasets lies in enabling trajectory-aware representations. More broadly, these findings challenge the common assumption that increasing feature richness through NLP necessarily improves predictive performance, highlighting instead the importance of representation design in low-sample, high-dimensional settings.

A first point that should be emphasized is the interpretation of the metric drop observed after leakage correction. This reduction should not be framed as a deterioration of the study, but rather as a methodological validation step. Data leakage is a well-recognized source of performance inflation in biomedical machine learning, particularly when preprocessing, feature selection, or other data-dependent transformations are applied before train-test separation [27,28]. The problem is especially relevant in small datasets, where performance estimates are intrinsically more variable and more sensitive to optimistic evaluation strategies [29,30]. In our case, the leakage-free pipeline produced more conservative metrics than earlier analyses, but these estimates are more likely to reflect the true generalization ability of the models. This interpretation is further reinforced by the broader literature on high-dimensional biomedical prediction, where unstable or inflated performance is common when sample size is limited relative to the number of features [29,30].

The second main result concerns the original hypothesis that free-text information from T1 would improve classification performance. Our results do not support that hypothesis in a robust way. Across the final analyses, Pool2 showed at most marginal gains over Pool1, and the ablation analysis indicated that removing the compact text block did not consistently degrade performance. This finding should not be interpreted as evidence that the text contained no clinically relevant information, but rather that its added predictive value was either redundant or too weak to be exploited reliably in the present setting. Two explanations seem particularly plausible. First, the information spontaneously reported by participants in free-text responses may already have been captured, at least partially, by the structured questionnaire variables. Second, in a cohort of approximately 100 participants, the available examples may simply be insufficient for stable extraction of useful static patterns from clinical text-derived variables. Both interpretations are consistent with the clinical NLP literature, which repeatedly highlights restricted data availability, single-site provenance, and overfitting risk as major constraints for text-based machine learning in healthcare [6,9,17].

The third and most relevant finding is that model performance improved only when the follow-up information was represented as compact longitudinal descriptors. This was the point at which Pool3 clearly separated itself from Pool1 and Pool2. Importantly, the gain was not obtained by merely adding more variables, but by changing the representational strategy. Instead of introducing raw T2 variables as another large feature block, the final pipeline summarized how patients changed between T1 and T2 through compact temporal descriptors. This distinction is critical. Previous methodological work has shown that temporal and longitudinal representations can provide a clinically meaningful predictive signal when they are formulated in a way that captures trajectories rather than static snapshots [11,31]. Our results are consistent with that idea: the best Pool3 model was obtained only after T2 was encoded as change, suggesting that the informative signal resided in the longitudinal pattern itself rather than in the isolated follow-up measurements.

The ablation analysis strengthens this interpretation. In Pool2, removing the text block did not produce a coherent loss of performance, indicating that static text-derived variables were not a stable driver of classification. In Pool3, removing the compact text block had only minimal impact overall, whereas removing the compact temporal block caused a marked and consistent decrease in performance across both holdout and cross-validation. This provides stronger evidence than simple between-pool comparison because it isolates the contribution of each block within the same modeling framework. Therefore, the principal contribution of the final pipeline does not seem to be “NLP enrichment” in the broad sense, but rather the ability to transform questionnaire information into a clinically meaningful summary of longitudinal change. In this respect, the study moves from asking whether text improves classification to identifying where language-based processing is actually useful in small clinical cohorts.

An additional point of interest is the behavior of the models themselves. In the final configuration, Random Forest showed the greatest improvement in Pool3. Although this should be interpreted cautiously, it suggests that the signal carried by the longitudinal descriptors may be at least partly nonlinear or interaction-dependent. Linear models, which were competitive in Pool1 and Pool2, may have been less able to exploit that structure. This interpretation is consistent with the idea that change-related variables may encode conditional or interacting relationships that are not fully captured by additive linear effects. Even so, our results do not imply that Random Forest is universally preferable; rather, they indicate that the choice of model depends on the structure of the representation being learned.

These findings also help refine the role of LLM-based feature construction in this context. The present results do not support the use of LLM-derived variables merely as a way to increase the number of static features in small cohorts. Under that framing, the gain was limited or inconsistent. However, the results do support a different and more defensible use case: employing language-based processing as a tool for abstraction and summarization, particularly when the goal is to encode temporal evolution in a compact and informative way. This distinction is important for future work because it suggests that the added value of NLP in clinical machine learning may depend less on semantic richness alone and more on how textual or semi-structured information is transformed into trajectory-aware representations.

Several limitations should be acknowledged. First, the sample size remained modest, which constrains both statistical stability and model complexity. Second, the study relied on repeated internal validation rather than external validation, so the generalizability of the final model to other cohorts remains to be established. The generalizability of these findings to larger cohorts or different disease contexts remains an open question and should be addressed in future work. Third, the derived variables depended on questionnaire content and on the chosen representation strategy; alternative temporal encodings could potentially perform differently. Fourth, the outcome remained a binary ALS-versus-control classification, meaning that the present study cannot yet address more granular phenotypic distinctions such as DFT or ALS-DFT. These limitations, however, do not weaken the central message of the work. On the contrary, they help define the setting in which the conclusions should be interpreted: a small clinical cohort, moderate class imbalance, and questionnaire-based multimodal information.

Taken together, the study supports a more precise framing than the one initially proposed. The results do not show that free-text processing, by itself, improves ALS classification. Rather, they indicate that the main value of language-based processing in this setting lies in its capacity to structure and summarize longitudinal change. This is the most important contribution of the work: not simply evaluating whether LLM-derived features help, but identifying under which representational conditions they become useful. Future studies should test whether this conclusion remains valid in larger cohorts, under external validation, and in settings where richer longitudinal text or more frequent follow-up assessments are available.

## 6. Conclusions

This study shows that the predictive value of questionnaire-derived information depends strongly on how that information is represented. After correcting information leakage and rebuilding the pipeline under a fully validation-aware framework, performance estimates became more conservative but also more credible, reinforcing the methodological validity of the study [27-30]. Under this leakage-free setting, variables derived from first-time-point free-text responses did not provide a robust and reproducible improvement over structured baseline variables alone.

The main contribution of the work emerged only after follow-up information was reformulated as compact descriptors of longitudinal change between T1 and T2. In this final representation, Pool3 consistently outperformed both Pool1 and Pool2, and the ablation analysis showed that removing the temporal block substantially degraded performance, whereas removing the compact text block had little or no negative effect. These findings indicate that the main source of improvement was not the inclusion of additional static variables, but the introduction of a more informative longitudinal representation.

Therefore, the results do not support a general claim that NLP-derived variables improve ALS classification by themselves. Instead, they support a more precise conclusion: in small clinical cohorts, the main value of language-based processing may lie in its ability to structure and summarize longitudinal change, rather than to enrich static feature spaces [11,31]. In this sense, the study identifies not only where LLM-based processing fails to add value, but also where it can become genuinely useful. This work suggests a shift in how NLP should be used in clinical machine learning: not as a tool for feature expansion, but as a mechanism for constructing meaningful representations of patient trajectories.

From a methodological perspective, this work highlights the importance of leakage-free evaluation, dimensionality control, and careful feature representation in questionnaire-based clinical prediction [11,27-31]. From an applied perspective, it suggests that future work should prioritize trajectory-aware representations, larger cohorts, and external validation settings in order to determine whether the same pattern holds beyond the present dataset. More broadly, the findings support the use of NLP and LLM-based approaches as tools for abstraction and temporal synthesis, rather than simply as generators of additional static features.

## List of abbreviations

ALS: Amyotrophic lateral sclerosis
ALS-FTD: Amyotrophic lateral sclerosis-frontotemporal dementia
ALS-FTSD: Amyotrophic lateral sclerosis-frontotemporal spectrum disorder
API: Application programming interface
BMI: Body mass index
CV: Cross-validation
EHR: Electronic health record
FTD: Frontotemporal dementia
IRB: Institutional Review Board
JSON: JavaScript Object Notation
LLM: Large language model
MCC: Matthews correlation coefficient
ML: Machine learning
NLP: Natural language processing
RF: Random Forest
SDoH: Social determinants of health
SVC: Support Vector Classification
T1: Time point 1
T2: Time point 2
WHO: World Health Organization

